# A Novel eDNA-Based Approach for Hybrid Detection: Implications for Conservation Management

**DOI:** 10.64898/2026.03.26.714632

**Authors:** Masayuki K. Sakata, Nanako Yano, Akio Imamura, Hiroki Yamanaka, Toshifumi Minamoto

## Abstract

Hybridization between invasive and native species poses a hidden but critical threat to biodiversity. While environmental DNA (eDNA) has revolutionized species monitoring, it has lacked the resolution to detect hybrid individuals. Here, we present the first experimental demonstration of hybrid identification using eDNA. Our method isolates a single cell in the environment (hereafter, eCell) and enables cellular-level analysis using multiplex digital PCR targeting nuclear markers from both parental species. Validation with controlled tank experiments using *Oncorhynchus masou masou* × *Salvelinus leucomaenis leucomaenis* hybrid individuals confirmed the method’s ability to separately detect hybrid individuals from co-habiting purebred parent individuals. This eCell analysis overcomes the limitations of traditional eDNA methods and offers a scalable, non-invasive tool for detecting cryptic hybridization. By enabling early and accurate detection of hybrid individuals, it supports timely conservation decisions, including management prioritization and the protection of purebred populations. This novel technique bridges a critical gap in conservation genetics and enhances eDNA’s utility for biodiversity management in the face of global change.

## Introduction

Invasive alien species represent one of the most pervasive and accelerating threats to global biodiversity, altering native ecosystems through mechanisms such as competition, predation, disease transmission, and hybridization with native taxa (Simberloff 1996; Theodoropoulos et al. 2025). Among these, hybridization poses a particularly insidious challenge. Unlike overt ecological impacts, hybridization often proceeds cryptically—without apparent morphological changes—until native genetic lineages are diluted or lost entirely (Todesco et al. 2016; Chan et al. 2019). This process, known as cryptic introgression, involves gradual genomic replacement without obvious morphological change and undermines species-based conservation by eroding endemic genotypes, blurring taxonomic boundaries and complicating the legal and ethical treatment of hybrid individuals (Jackiw et al. 2015).

Human-mediated environmental changes are increasingly intensifying hybridization risks. The introduction and translocation of non-native species, together with anthropogenic habitat alteration, have promoted contact between previously isolated taxa, often leading to unexpected or asymmetric hybridization events (Muhlfeld et al. 2014; Bohling 2016; Ottenburghs 2021). Such human-facilitated interactions can result in so-called “extinction without extinction,” where species remain phenotypically recognizable but lose their evolutionary distinctiveness through genome-wide introgression (Chan et al. 2019; Fitzpatrick et al. 2024). While hybridization can occasionally promote adaptive evolution (vonHoldt et al. 2017), current conservation frameworks are ill-equipped to assess or mitigate these complex outcomes (Wayne & Shaffer 2016). Even when hybridization is identified as a conservation threat, management efforts often target only the parental species, overlooking hybrids themselves (Zbinden et al. 2023). This challenge is compounded by the lack of sensitive and field-deployable tools to detect early-stage or cryptic hybridization in natural populations (Kuijk et al. 2025; Rosas et al. 2025).

Early detection of non-native species and hybrid individuals is crucial for biodiversity conservation because it allows for rapid management responses before irreversible genetic or ecological impacts occur. Environmental DNA (eDNA) analysis has become a powerful and widely adopted approach for detecting invasive species, owing to its high sensitivity and scalability across large spatial extents (Goldberg et al. 2016; Minamoto 2022; Takahashi et al. 2023). eDNA has been applied to species distribution surveys, biomass estimation, and even population genetic inferences in invasive species contexts (Sepulveda et al. 2020; Rourke et al. 2021; Yatsuyanagi et al. 2024). However, conventional eDNA techniques extract bulk DNA from environmental samples, which consist of both extracellular and intracellular DNA originating from multiple individuals. Consequently, they cannot provide information at the individual level and are unable to detect hybrid individuals (Minamoto 2022). To date, no published studies have successfully identified hybrid individuals using eDNA-based methods. To overcome this limitation, a methodological breakthrough is needed—specifically, a way to detect nuclear markers derived from both parental species within a single cell in the environment (hereafter, eCell). If hybrid cells can be isolated and analyzed without destroying cellular integrity, it becomes possible to identify hybrid individuals directly from environmental samples. In this study, we explore this possibility by developing and applying a novel framework that independently analyzes two parental nuclear markers within each eCell using digital PCR (dPCR) to determine the presence of hybrid individuals.

Here, we aimed to develop and validate an eCell approach to detect hybrid individuals, with direct implications for conservation management. We validated this method through a two-step process. First, we confirmed that dPCR enables the simultaneous co-detection of two genomic markers within a single cell isolated from tissue samples. Second, we then applied the technique to experimental tanks containing hybrid individuals derived from two salmonid fish species, *Oncorhynchus masou masou* × *Salvelinus leucomaenis leucomaenis*, and these salmonid species produce sterile F□ individuals. We demonstrate the first feasible approach for detecting the presence of hybrid individuals from environmental samples, with potential applications for timely conservation interventions. This approach provides a scalable and field-adaptable framework for detecting hybridization and monitoring genetic introgression. While further validation in natural settings is needed, the method holds strong potential to inform timely conservation interventions, such as the removal of hybrid individuals or the prioritization of management zones, before irreversible genomic erosion occurs.

## Materials and Methods

### Confirmation of cell-by-cell DNA dPCR

We performed multiplex dPCR targeting mitochondrial DNA (mtDNA) and nuclear DNA (nuDNA) of *Hemigrammocypris neglecta* to confirm the simultaneous co-detection of both markers within a single cell isolated from tissue samples. We hypothesized that, because each dPCR reaction contains a large number of partitioned wells, the probability of two different DNAs co-occurring in a single well is very low for extracellular DNA, whereas it should be substantially higher for single cell (i.e., within individual cells). Accordingly, the number of wells in which two markers (e.g., mtDNA and nuDNA, or nuclear markers from both parental species) are simultaneously detected is expected to exceed the range predicted by random co-detection of extracellular DNA. To test this, we simulated the expected distribution of co-detection under the assumption that all DNA is extracellular and fully dissociated, so that mitochondrial DNA, nuclear DNA, and even DNA from different chromosomes is present as separate fragments. We then compared this with the observed number of wells containing two markers simultaneously, thereby evaluating the potential of dPCR for single-cell DNA detection (Fig. 1). For details on the simulation, see the Supplementary Information.

**Fig. 1.**
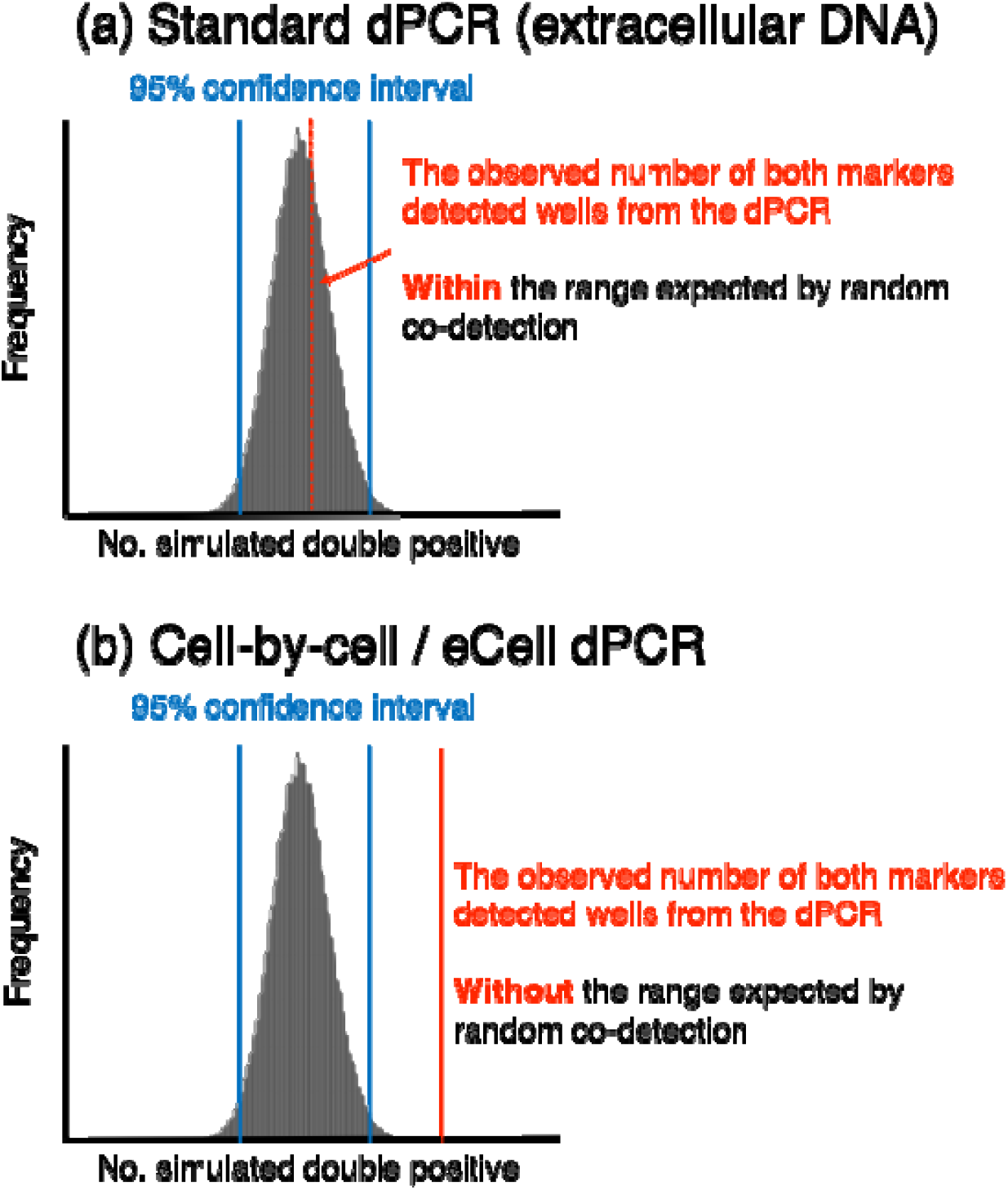
Conceptual diagram of hypothesis verification. The blue line shows the 95% confidence interval of the simulation assuming random distribution of two marker genes. The red dotted line shows the observed number of both markers detected wells from the dPCR. The x-axis shows the simulated number of dPCR wells in which two marker genes are co-detected. The y-axis indicates the frequency of occurrence. (a) Cases in which the observed number of wells detecting both markers falls within the range expected by random co-detection. In this scenario, cell-by-cell dPCR is considered not established. (b) Cases in which the observed number of wells detecting both markers exceeds the range expected by random co-detection. In this scenario, cell-by-cell dPCR is considered successfully established.

Genomic DNA was extracted from the muscle tissue using the DNeasy Blood & Tissue Kit (QIAGEN, Hilden, Germany) following the manufacturer’s protocol. Single cells were isolated from the muscle tissue of *H. neglecta* using the 10-min Tissue (Tumor) Dissociation / Single Cell Isolation Kit (101 Bio, LLC) according to manufacturer instructions. The dPCR assays were conducted in duplicate using either extracted genomic DNA or isolated single cell suspension as templates. The isolated single-cell suspension was prepared by suspending cells isolated from tissue samples in 1× PBS buffer. Reactions were performed on the QIAcuity One Platform System (QIAGEN) using QIAcuity Nanoplate microfluidic dPCR plates, which allow processing of up to 24 samples with up to 26,000 wells per reaction. Each 40 µL reaction contained 800 nM primers (Hra-cytb-F/R for mtDNA and Hra-ITS1-F/R for nuDNA), 400 nM TaqMan probes (Hra-cytb-P and Hra-ITS1-P), 1 × QIAcuity OneStep Advanced Probe MasterMix (QIAGEN), and 3 µL of template DNA or isolated cell suspension. The Primers and probe were developed in previous studies (Fukuoka et al. 2016; Hashizume unpublished) and those sequences are listed in Table 1. Thermal cycling conditions were: initial denaturation at 95 °C for 2 min, followed by 45 two-step cycles of 15 s at 95 °C and 30 s at 60 °C.

**Table 1:**
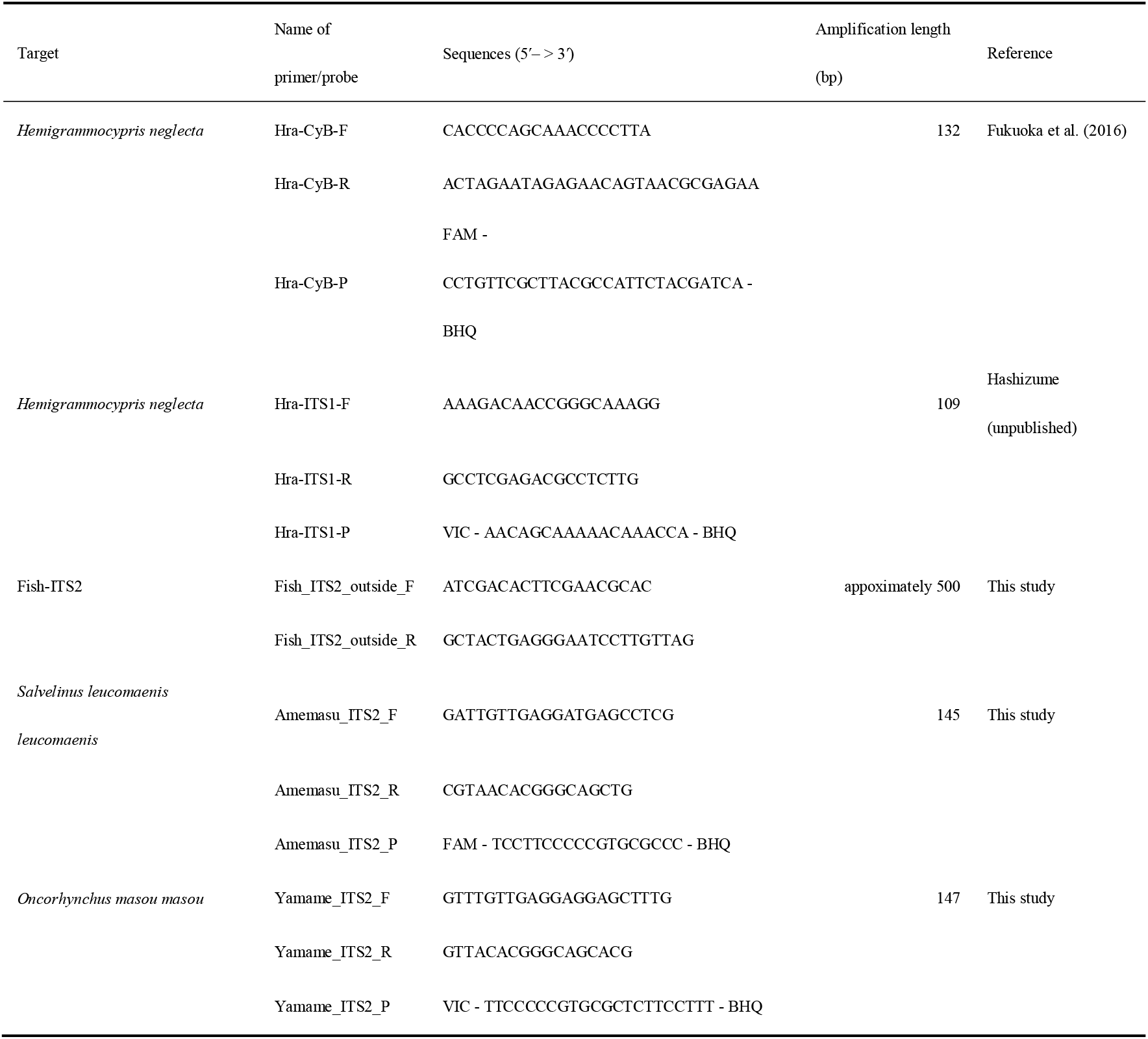
List of primers and probes used in this study.

To evaluate co-detection of mtDNA and nuDNA within the same wells, we simulated the expected number of wells positive for both targets based on observed counts of wells positive for mtDNA, nuDNA and total wells. The simulation was repeated 100,000 times using the *sample* function in R (version 4.5.1), calculating the 95% confidence interval. The actual number of wells positive for both mtDNA and nuDNA in dPCR experiments was then compared to this interval to assess whether the observed co-detection exceeded random expectation, thus validating the feasibility of cell-by-cell dPCR. Analysis scripts are provided in the Supplementary Information. All graphs were generated with the ggplot2 package (v3.5.2).

### Development and validation of the dPCR assay

New primers and probes were designed targeting nuclear DNA regions specific to *O. m. masou* and *S. l. leucomaenis*. DNA was extracted from three *O. m. masou* and four *S. l. leucomaenis* individuals using the DNeasy Blood & Tissue Kit (QIAGEN) per manufacturer’s instructions. Approximately 500 bp fragments were amplified using primers Fish_ITS2_outside_F and Fish_ITS2_outside_R (Table 1) with TaKaRa Ex Taq Hot Start Version (Takara Bio, Shiga, Japan). Each 25 µL PCR reaction contained 2.5 µL 10× Ex Taq Buffer, 2 µL dNTPs (2.5 mM each), 400 nM of each primer, 0.125 µL TaKaRa Ex Taq HS, and 2 µL template DNA. Cycling conditions were initial denaturation at 94 °C for 2 min, followed by 30 cycles of 10 s at 98 °C, 30 s at 55 °C, and 60 s at 72 °C. PCR products were purified using the Wizard SV Gel and PCR Clean-up System (Promega) and commercially sequenced by FASMAC (Tokyo, Japan) with the same primers. We obtained the three *O. m. masou* ITS2 sequences and four *S. l. leucomaenis* ITS2 sequences (LC920519, LC920520, LC920521, LC920522, LC920523 LC920524 and LC920525). Based on those sequences, primers (Yamame_ITS2_F, Yamame_ITS2_R, Amemasu_ITS2_F and Amemasu_ITS2_R) and probes Yamame_ITS2_P and Amemasu_ITS2_P were designed (sequences in Table 1). Based on obtained sequences and following Sakata et al. (2017), species-specific primers and TaqMan probes were designed for *O. m. masou* and *S. l. leucomaenis*. Specificity and efficiency of assays were confirmed by dPCR in duplicate using eCell samples (see below) of each species on the QIAcuity platform. Each 40 µL reaction contained 400 nM primers (Amemasu_ITS2_F/R for *S. l. leucomaenis* and Yamame_ITS2_F/R for *O. m. masou*), 200 nM TaqMan probes (Amemasu_ITS2_P and Yamame_ITS2_P), 1 × QIAcuity OneStep Advanced Probe MasterMix (QIAGEN), and 2 µL template DNA (100 pg per reaction) from tissue-derived DNA of parental species and hybrid individuals. Thermal cycling was 2 min at 95 °C followed by 45 two-step cycles of 15 s at 95 °C and 30 s at 62 °C. Primer and probe sequences are in Table 1.

### Validation of hybrid individuals detection using eCell analysis

Tank experiments were conducted at Kobe University from 26 March to 16 April 2024 to validate hybrid detection using intracellular analysis. Tanks contained either two hybrid individuals (*O. m. masou* × *S. l. leucomaenis*) or one individual of each parental species, caught in Hokkaido, Japan. Two fish were housed per tank with 10 L of water. To collect eCell samples, 6 L of water from each tank was immediately filtered through a 30 µm nylon filter (Merck Millipore, Inc.). eCell samples trapped by the filters were resuspended in 5 mL of 1× PBS buffer and kept on ice until dPCR. To prevent cross-contamination, all equipment used for water collection and filtration (plastic bottles, funnels, tweezers) were decontaminated with >0.1% sodium hypochlorite solution (Minamoto et al. 2021). dPCR was performed in duplicate on eCell samples using the same platform and conditions as described in “Development and validation of the dPCR assay.” Non-template controls (NTCs) with ultrapure water instead of DNA were included to detect contamination. Sampling and dPCR were repeated four times: twice for hybrid tanks and twice for parental tanks. Each dPCR result was evaluated via the same random simulation approach as in “Confirmation of cell-by-cell dPCR” to determine whether the number of wells simultaneously positive for nuclear DNA of *O. m. masou* and *S. l. leucomaenis* exceeded random expectation.

## Results

### Confirmation of cell-by-cell dPCR

The simulation results showed that, for the gDNA sample, the observed number of wells in which both markers were detected fell within the predicted range (95% confidence interval) assuming they were randomly distributed to the wells (Fig. 2a). In contrast, for the isolated single cell sample, the observed number of wells in which both markers were detected exceeded the 95% confidence interval predicted by the random distribution (Fig. 2b).

**Fig. 2.**
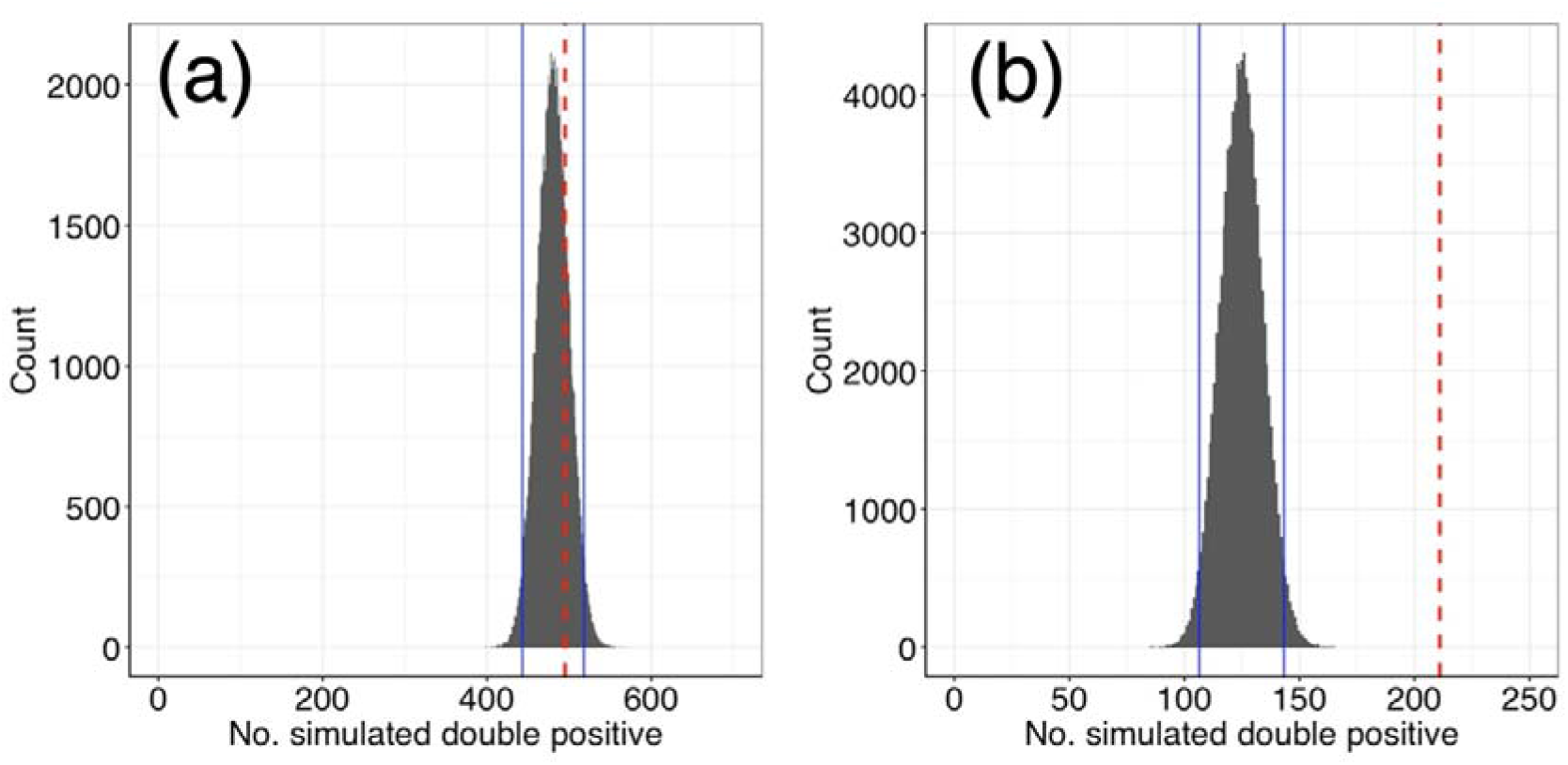
The results of dPCR using tissue derived gDNA and isolated cell sample of *Hemigrammocypris neglecta*. The blue line shows the 95% confidence interval of the simulation assuming random distribution of two marker genes. The red dotted line shows the observed number of both markers detected wells from the dPCR. (a) The genomic DNA sample from tissue. The numbers of wells were as follows: 9,996 positives only for mtDNA, 1,838 only for nuDNA, 495 for both, and 38,607 negatives. (b) The isolated cell sample from tissue. The numbers of wells were as follows: 14,573 positives only for mtDNA, 219 only for nuDNA, 211 for both, and 35,899 negatives. The observed number of both markers detected wells fell within the range of randomness for (a), but not for (b).

### Validation of hybrid individuals detection using eCell analysis

In the specificity and effectiveness test by dPCR, each assay amplified the target species DNA only. In addition, the DNA derived from a hybrid individual was positive in both assays. No amplification was detected in any NTC in either assay.

In the dPCR results from eCell samples obtained from a tank containing one individual each of *O. m. masou* and *S. l. leucomaenis*, both markers of those target species were amplified. The number of wells in which both markers were simultaneously detected was within the predicted range of random distribution in both of two trials (Fig. 3a, b). In contrast, in the dPCR results from eCell samples obtained from a tank containing only two hybrid individuals, both markers of those target species were amplified, and the number of wells in which both markers were simultaneously detected exceeded the predicted range of randomness (Fig. 3c, d).

**Fig. 3.**
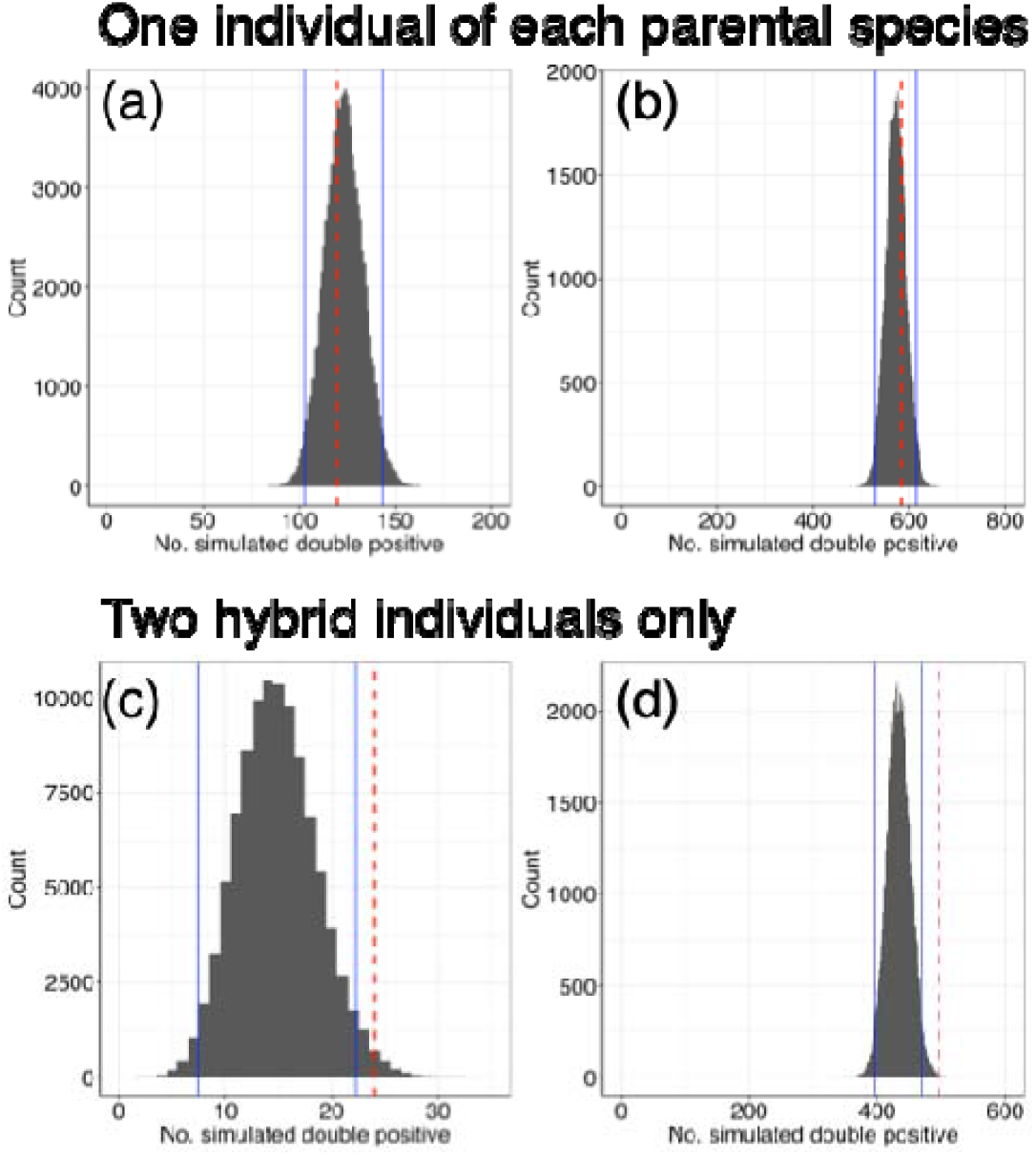
The results of dPCR by eCell samples from each tank experiment (parental species individuals tanks and hybrid individuals only tanks). The blue line shows the 95% confidence interval of the simulation assuming random distribution of two marker genes. The red dotted line shows the observed number of both markers detected wells from the dPCR. (a) and (b) The dPCR and simulation results for a sample of aquarium water containing one individual of each parent species. The numbers of wells were as follows: 7,411 and 6,455 positives only for *O. m masou*, 711 and 3,545 only for *S. l. leucomaenis*, 120 and 582 for both, and 42,550 and 40,251 negatives, respectively. (c) and (d) The dPCR and simulation results of a sample of aquarium water containing two hybrid individuals. The numbers of wells were as follows: 631 and 3,050 positive only for *O. m. masou*, 1,131 and 5,662 only for *S. l leucomaenis*, 24 and 495 for both, and 49,122 and 41,210 negative, respectively. The observed number of both markers detected wells fell within the range of randomness for (a) and (b), but not for (c) and (d).

## Discussion

This study demonstrates for the first time that hybrid individuals can be identified using eDNA and advances the field by extending eDNA applications to individual-level conservation genetics. The results of the two types of tank experiments clearly demonstrate that our developed method can distinguish between the co-habiting of parental species and the presence of hybrid individuals, thereby enabling the detection of hybridization. This novel analytical method was established by recovering eCell and analyzing it without disrupting cell integrity using dPCR to simultaneously detect genetic markers of two parental species from a single cell as a proof of hybridization. Applying this novel approach to field studies will not only allow for the rapid detection of invasive species but also determine whether hybridization with native taxa is occurring. This substantially enhances one of the primary strengths of eDNA analysis, namely its ability to conduct rapid, large-scale monitoring. Previous eDNA-based studies on invasive species have focused on estimating species distributions, biomass, or community-level impacts (Dougherty et al. 2016; Takaba et al. 2024; Wakimura et al. 2025). Our method addresses this gap, enabling the detection of hybridization and advancing the use of eDNA in managing biological invasions. Importantly, by revealing the occurrence of hybridization in invaded areas, this method provides essential information for invasive species management. It supports the preservation of genetic diversity, the identification of critical zones where immediate action is required, and the prioritization of areas for conservation and control efforts. As such, it offers a powerful tool for guiding evidence-based decision-making in biodiversity conservation. The need for conservation actions grounded in systematic evidence rather than anecdotal practice has been repeatedly emphasized in the conservation literature (Sutherland et al. 2004). By providing direct genetic evidence of hybridization at the individual level, our framework contributes to closing this implementation gap.

Hybridization is often overlooked in conservation management due to the difficulty of detecting it in its early stages, especially when hybrid individuals are morphologically indistinguishable or occur at low densities (Zbinden et al. 2023; Kuijk et al. 2025; Rosas et al. 2025). Our novel eDNA-based method has the potential to overcome these challenges. Unlike traditional methods, eDNA detection does not require capturing individuals or relying on morphological traits for identification (Minamoto 2022). Moreover, eDNA sampling can reflect biological presence over spatial ranges of several meters to several hundred meters, even when target organisms occur at low densities (Barnes & Turner 2015; Jo & Yamanaka 2022). These features make eDNA analysis particularly well-suited to detecting hybridization at early or cryptic stages. Early and sensitive detection is critical because once introgression spreads within populations, management becomes increasingly difficult, costly, and often irreversible (Todesco et al. 2016). As hybrid individuals accumulate and genetic swamping progresses, eradication or restoration of purebred populations may become impractical (Todesco et al. 2016). Such genome-level replacement, often termed genetic swamping, can lead to the extinction of pure parental genotypes even when phenotypic forms persist (Wayne & Shaffer 2016). This process complicates conservation policy, as legal frameworks are typically designed to protect species rather than genomes or specific genetic lineages (Allendorf et al. 2001). Recent perspectives emphasize the need to evaluate hybridization at the genic level, recognizing that introgression can differentially affect genomic regions and conservation value (Ottenburghs 2021). By enabling accurate and rapid detection of hybridization at its earliest stages, our technology supports timely intervention before ecological and genetic impacts escalate. It allows managers to detect hybridization before introgression exceeds recoverable levels, enabling intervention at ecologically meaningful thresholds. Defining such thresholds remains a central challenge in hybrid management, as conservation decisions often depend on whether genetically pure individuals persist or whether hybrid swarms have formed (Allendorf et al. 2001). Early detection is therefore critical to prevent irreversible genomic replacement (Wayne & Shaffer 2016). Furthermore, eDNA-based detection provides direct information for conservation action, including the identification of priority management zones and the protection of genetically pure populations (Bohling 2016). A unique advantage of this approach is its ability to simultaneously detect both parental species and hybrid individuals from the same environmental sample, thereby enhancing monitoring efficiency and providing richer genetic context. Taken together, our eCell framework offers a scalable and decision-relevant tool for conservation systems where hybridization poses a critical risk.

While this study provides the first proof-of-concept for detecting hybrid individuals using eDNA-based cell isolation and eCell dPCR, further validation under natural field conditions is essential before practical application. Environmental complexities—such as DNA degradation, microbial activity, and the presence of mixed DNA from multiple individuals—may influence the sensitivity and specificity of hybrid detection. To address these challenges, future work should test this approach in natural systems where hybridization dynamics and population structures are already well characterized. Establishing the robustness of this method in real-world settings will be critical for its integration into conservation practice. Once validated, this approach can be integrated into genetic monitoring programs to proactively manage hybrid zones, protect genetically pure populations, and prevent irreversible genetic erosion. By enabling genome-informed detection of hybridization at the level of individual cells, our framework operationalizes recent calls to integrate conservation genomics into management under accelerating anthropogenic hybridization (Ottenburghs 2021). As hybridization increasingly shapes biodiversity in the Anthropocene (Ottenburghs 2021)., tools that distinguish between genomic erosion and adaptive introgression will be central to maintaining evolutionary integrity.

## Acknowledgement

We are grateful to I. T. Hirayama and Hokkaido Mizubenokai TK for their support to conduct the tank experiment and for the sample collection, respectively.

