## Supplementary material for "A Novel eDNA-Based Approach for Hybrid Detection: Implications for Conservation Management": Supplemntal Information

Supplementary Information

#no: the number of no detection wells

#nu: the number of wells detected in one of the two markers

#mt: the number of wells detected in the other marker

\# real_both: the number of wells detected in both markers

#n: the number of iterations

#X_upper: the upper limit of the x-axis

function_dPCR <- function(no, nu, mt, real_both, n, X_upper) {

total <- mt + nu + real_both + no

mito <- mt + real_both

nucl <- nu + real_both

wells <- matrix(0, total, 2)

colnames(wells) <- c("mt", "nu")

wells[1:mito, 1] <- 1

wells[1:nucl, 2] <- 1

result <- c()

for (i in 1:n) {

re_mt <- sample(wells[,1])

re_nu <- sample(wells[,2])

wells_random <- data.frame(re_mt, re_nu)

both <- rowSums(wells_random)

table(both)

Wpositive <- sum(as.integer(both > 1))

result <- rbind(result, c(i, Wpositive))

}

result

mean <- mean(result[, 2])

sd <- sd(result[, 2])

conf_u <- mean + 1.96*sd

conf_l <- mean - 1.96*sd

g <- ggplot() +

geom_histogram(aes(result[,2]), binwidth = 1) +

theme_bw() +scale_x_continuous(limits = c(0, X_upper))+

labs(x = 'No. simulated double positive', y = 'Count')+

geom_vline(xintercept = real_both, colour = "red", size = 1, linetype = 2) +

geom_vline(xintercept = conf_u, colour = "blue", size = 0.5, linetype = 1) +

geom_vline(xintercept = conf_l, colour = "blue", size = 0.5, linetype = 1) +

theme(

axis.title = element_text(size = 20),

axis.text = element_text(size = 20,colour = "black"),

text = element_text(family = 'Helvetica'),

legend.position = 'right',

axis.line=element_line(colour = "black"),

axis.ticks=element_line(colour = "black")

)

g

}
